# Reinforced molecular dynamics: Physics-infused generative machine learning model explores CRBN activation process

**DOI:** 10.1101/2025.02.12.638002

**Authors:** István Kolossváry, Rory Coffey

**Affiliations:** Biokol Research Consulting, Acton, Massachusetts, U.S.A; Rothco Science Solutions, Boston, Massachusetts, U.S.A

## Abstract

We propose a simple and practical machine learning-based desktop solution for modeling biologically relevant protein motions. We termed our technology reinforced molecular dynamics (rMD) combining MD trajectory data and free-energy (FE) map data to train a dual-loss function autoencoder network that can explore conformational space more efficiently than the underlying MD simulation. The key insight of rMD is that it effectively replaces the latent space with an FE map, thus infusing the autoencoder network with a physical context. The FE map is computed from an MD simulation over a low-dimensional collective variable space that captures some biological function. One can directly use then the FE map for example, to generate more protein structures in poorly sampled regions, follow paths on the FE map to explore conformational transitions, etc. The rMD technology is entirely self-contained, does not rely on any pre-trained model, and can be run on a single GPU desktop computer. We present our rMD computations in a key area of molecular-glue degraders aimed at a deeper understanding of the structural transition from open to closed conformations of CRBN.

---

Large machine learning (ML) systems AlphaFold (*1, 2*), RosettaFold (*3*), OpenFold (*4*), and ESM-Fold (*5*) revolutionized the field of protein structure prediction and spawned new research to produce ever more powerful computational models, e.g., (*6–8*). These systems are now widely available for the research community and have quickly become staples in the computational toolkit of the structural biology and drug discovery research fields. Notwithstanding their stunning success, these models predict static structures and cannot directly include protein motion, which defines most biological processes at the molecular level. In recent years, a great deal of work has been dedicated to combine these ”fold” algorithms with molecular dynamics (MD) simulations to inject dynamic data into ML models (*9–21*). At the heart of all of these ML systems lies the so-called autoencoder. The autoencoder can be visualized as a butterfly, with the thorax representing the so-called latent space. In our context, the autoencoder takes a protein structure as input, gradually compresses it into a low-dimensional latent space, and then expands it back to the real space. After training the autoencoder on large amounts of data from MD simulation trajectories, arbitrary points in the latent space can be reconstructed in real space, providing potentially interesting protein structures (*9*). The variational autoencoder design goes one step further by injecting additional physics-based information into the latent space (*11–14, 16*) in the form of a prior probability distribution with tunable parameters that are co-optimized with the network parameters during the training process in a Bayesian inference framework. The key idea behind our LM-reinforced molecular dynamics algorithm (rMD) is not to use a prior parametric distribution to tweak the latent space but to entirely replace it with a physical space, specifically a free energy (FE) map (*22*).

These advances in ML-driven structural modeling not only enhance our ability to predict protein conformations but also provide new insights into biologically relevant structural transitions. One such transition, central to targeted protein degradation, occurs in the E3 ligase substrate receptor, cereblon (CRBN).

Molecular-glue degraders are small molecules that facilitate a specific interaction between an E3 ligase and a target protein, leading to the proteolysis of the target. The discovery of molecular glue degraders targeting CRBN, an E3 substrate receptor protein, stemmed from the serendipitous identification of immunomodulatory imide drugs (IMiDs), a class of compounds first synthesized in 1954 (*23*). In 2010, thalidomide, one of these IMiDs, was revealed to bind to CRBN (*24*), and by 2014, this interaction was shown to induce the degradation of target proteins (*25*). This landmark finding, where IMiD binding directs the selective degradation of specific targets, has driven the emergence of a new pharmaceutical paradigm. This strategy has successfully enabled the targeted degradation of proteins such as GSPT1, IKZF1, IKZF3, CK1𝛼, VAV1, and NEK7 (*26–29*).

The development of IMiDs not only has expanded the range of degradable targets, but also has deepened our understanding of their effects on CRBN. Upon IMiD binding to CRBN, the protein undergoes a conformational shift from an inactive open state to an active closed state, as shown in Fig. 1 (*30–32*). However, this conformational transition remains underutilized in the discovery of new E3 degraders with similar functionality. A deeper understanding of the structural transition from open to closed states could facilitate the identification of novel E3 degraders and inspire the design of next-generation IMiDs with broader therapeutic applications.

**Figure 1:**
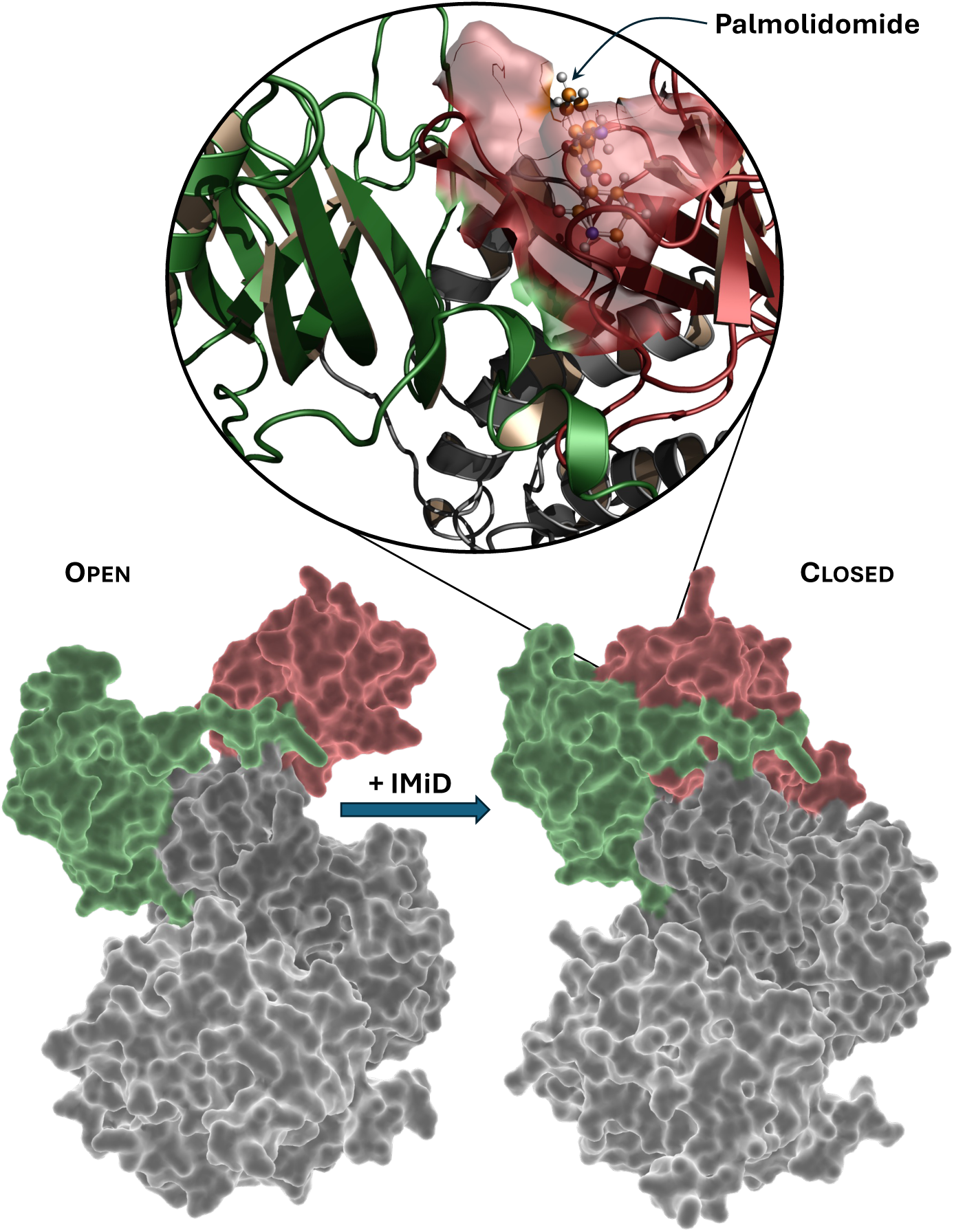
Conformational Transition of CRBN Upon IMiD Binding. CRBN undergoes a structural transition from an inactive open state to an active closed state upon IMiD binding. The structures shown are CRBN-DDB1 X-ray structures in the closed (PDB: 6H0G) and open (PDB: 6H0F) conformations. The CRBN Lon-like domain (LLD) is highlighted in green, and the thalidomide-binding domain (TBD) in red, emphasizing key structural changes induced by IMiD binding.

## Reinforced molecular dynamics

Reinforced molecular dynamics (rMD) is a comprehensive simulation technique that combines advanced sampling methods with a new kind of informed autoencoder model. Advanced sampling algorithms such as the many flavors of metadynamics (*33–39*) not only accelerate sampling along biologically relevant concerted atomic motions, termed collective variables (CV), but also provide the free-energy landscape associated with the CVs at hand. In (*22*) we present a detailed account of how we defined CVs to model CRBN closing and opening, and performed meta-eABF simulations (*22*) to generate a 3-dimensional free energy map associated with the chosen CVs. The key idea of rMD is to link this free energy map to the latent space of an autoencoder network.

The basic design of the autoencoder for protein structure prediction (*9*) is shown in Fig. 2 and a more detailed flow chart in fig. S2. The encoder passes the flattened Cartesian coordinates of protein structures saved during an MD simulation down a series of fully connected and gradually shrinking hidden layers. The final layer is called latent space (LS), denoted by a magenta-colored dot, where the original protein structures are compressed into a representation of only a few dimensions. The decoder then takes the low-dimensional latent space coordinates as input and decompresses them back to the original Cartesian representation using a similar set of now expanding hidden layers. As an example, we show two pairs of CRBN structures, the blue structures are inputs taken from the MD simulation, and the red structures are the network generated output structures. The basic autoencoder is trained on the ”predLoss” loss layer, minimizing the Loss2 loss function, which is the average root mean square distance (RMSD) between inputs and their predicted outputs; see more details in (*22*).

**Figure 2:**
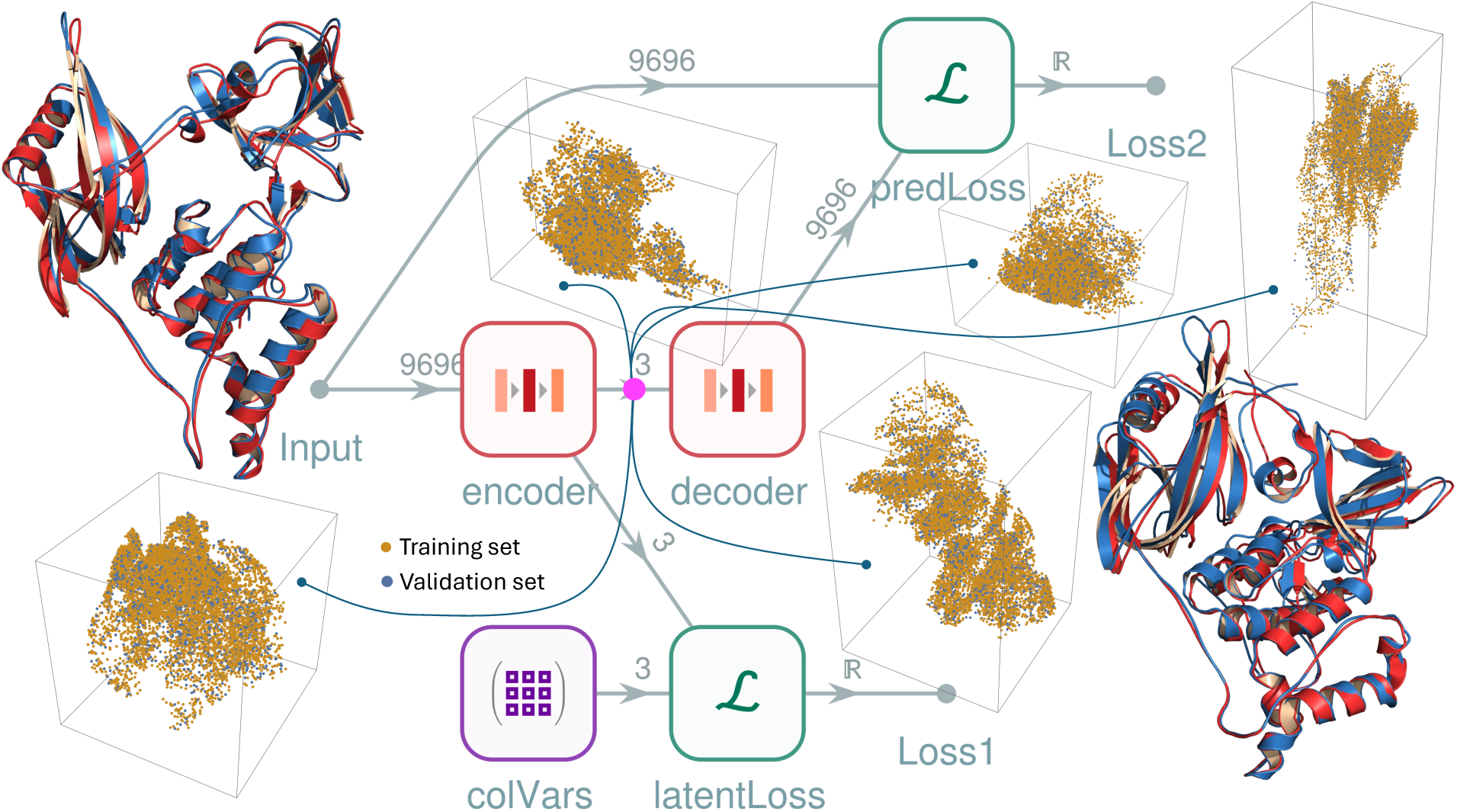
Network layout of the rMD informed autoencoder. The basic encoder compresses the flattened Cartesian coordinates of protein structures from MD simulations into a low-dimensional latent space (magenta dot) via fully connected network layers gradually shrinking in size. The decoder then reconstructs the original Cartesian representation from the LS using expanding layers. The blue CRBN structures represent inputs from simulations, while the red structures are their network-generated outputs. The autoencoder is trained using the ”predLoss” layer, minimizing the Loss2 function, which calculates the average RMSD between inputs and outputs. The 5 different point clouds show 5 different LS point distributions (orange-training set, blue-validation set) after training the basic network, initializing it with different random seeds. The informed autoencoder introduces an additional loss layer, ”latentLoss” and a loss function Loss1. The input for Loss1 are the LS coordinates and the target are the CV coordinates (purple ”colVars” box). By simultaneously optimizing Loss1 and Loss2, the trained network will compress the original trajectory frames into a unique latent space where LS coordinates have an approximate one-to-one correspondence with the CV coordinates as shown in Fig. 3. See main text and (*22*) for more details.

The basic autoencoder is a successful proof of concept for generating protein-like structures (*9*) but it is entirely generic and physics agnostic. To illustrate this, we trained the basic autoencoder 5 times, initializing it with 5 different random seeds. The different random seeds also resulted in 5 different randomized sets of corresponding training data (80%) and validation data (20%). In Fig. 2 we plotted 5 different point clouds representing the five trained distributions of the input data in the 3-dimensional latent space. The training set points are orange, and the validation points are blue. These vastly different latent space distributions vividly demonstrate a well-known crucial aspect of neural networks. The optimized network parameters represent a single local minimum among a plethora of other minima on an unimaginably complex ultra-high-dimensional surface, but experience shows that these minima are usually comparable in terms of the overall prediction power of the network (*40*). In our case, the five basic autoencoder networks predicted the CRBN structures with an average of ≈ 1.2 Å all heavy-atom RMSD. This is all good for generic structure generation, but does not work for targeted structure generation where we want to enlist machine learning (ML) techniques to find biologically relevant protein structures in a particular, e.g., drug discovery context. When we pick a point in the latent space, all we can expect is that the network will generate a protein-like structure. The basic tenet of rMD is to overcome this ambiguity in structure prediction by linking the latent space to a physical space of the same dimensions, such as the free energy map computed from an advanced sampling MD simulation.

The key idea to achieve this was to introduce an additional loss layer, ”latentLoss” and a loss function Loss1, as shown in Fig. 2 and fig. S2. Our informed autoencoder design empowers the network to perform targeted protein structure prediction by linking the latent space with the collective variable space. The input for the latent space loss function Loss1 are the LS coordinates and the target are the CV coordinates shown as the purple ”colVars” box. By simultaneously optimizing Loss1 and Loss2, e.g., using a weighted sum, the trained network will compress the original trajectory frames into a unique latent space where LS coordinates have an approximate one-to-one correspondence with the CV coordinates. This is how physics-based information is injected into the LS space. This is demonstrated in Fig. 3 where we plotted side by side the trained latent space points of the 10 000 trajectory frames on the left, and on the right we show a 3-dimensional density plot of the free energy map. The coloring represents a green-yellow-red color gradient from zero to 15 kcal/mol in relative free energy. The colors in the latent space were assigned by snapping the CV coordinates of each trajectory frame to the closest grid point in the free energy map. The huge difference in resolution between the two sides originates from only 10 000 points on the left and over 24 million points on the right (*22*). Nevertheless, the structural similarity between the latent space and the free-energy map in the CV space is striking, and it means that one can use the free-energy map directly to generate potentially interesting protein structures in low-free-energy (green) volumes.

**Figure 3:**
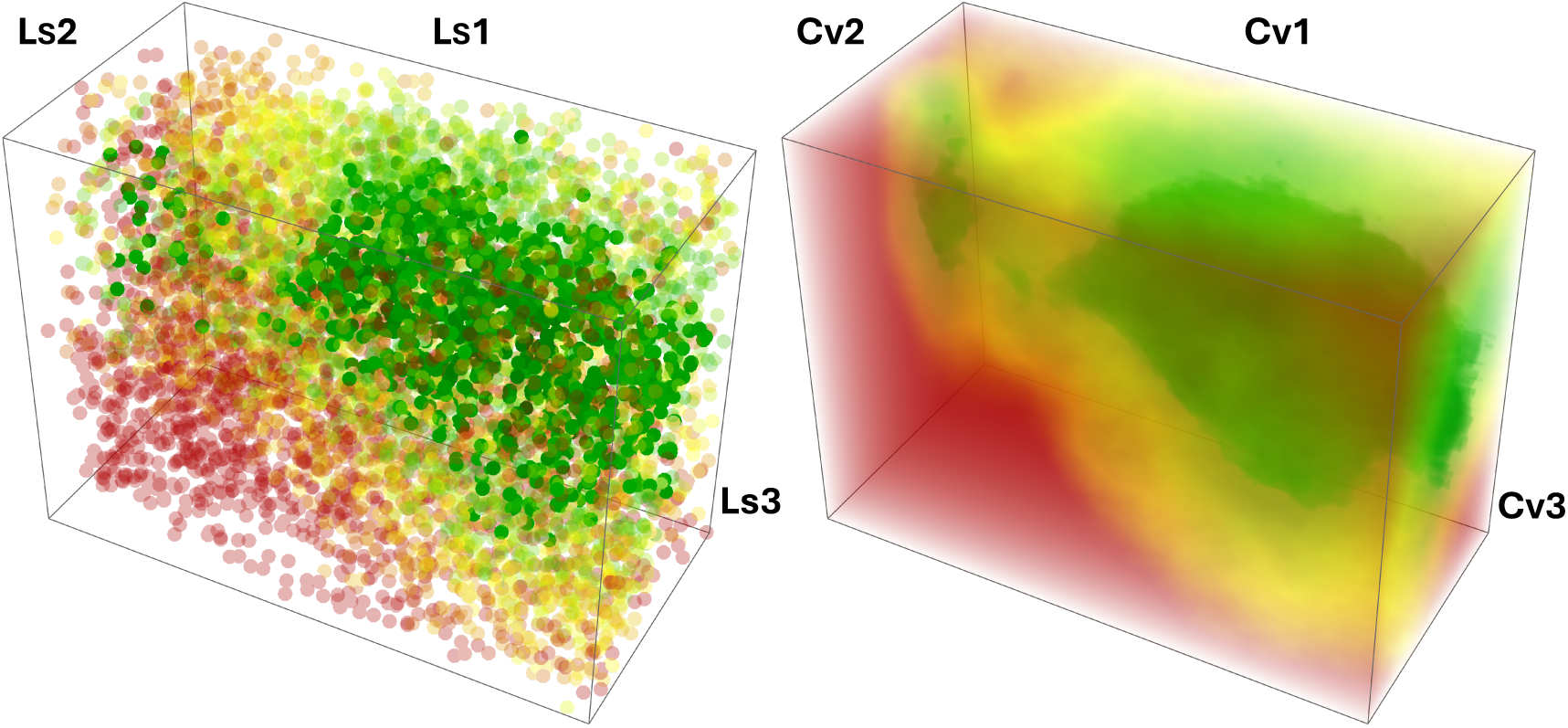
Visual comparison of the trained latent space of the rMD autoencoder and the free energy map in the collective variable space. We plotted side by side the trained latent space points of the 10 000 trajectory frames on the left, and on the right we show a 3-dimensional high-resolution density plot of the free energy map. The coloring represents a green-yellow-red color gradient from zero to 15 kcal/mol in relative free energy. The structural similarity between the latent space and the free-energy map in the CV space is striking, and it means that one can use the free-energy map directly to generate potentially interesting protein structures in low-free-energy (green) regions.

In Fig. 4 we demonstrate a typical use of rMD. The large black dot on the left shows the location of the X-ray structure of the closed CRBN conformation (PDB ID 6H0G) in the CV space, and the black dot on the right shows the location of the open CRBN conformation (PDB ID 6H0F) (*41*). The definition of CV coordinates is given in fig. S1. The free energy map shows that the volume of accessible closed conformations is very small compared to the very large number of feasible open conformations, and there is only a single narrow path connecting them. This makes perfect qualitative sense, because specific interactions between the CRBN-NTD (green) and CRBN-CTD (red) lobes are responsible for closing the conformation (*31, 32*), while free gyration of the center of mass of the CRBN-CTD lobe makes the open conformation very flexible, as shown in fig. S1. To establish a putative transition mechanism, we manually picked a few anchor points in the open, closed, and transition regions, respectively, then fitted a B-spline on them (blue curve), and computed the CV coordinates of the magenta points along the curve shown in Fig. 4 using the parametric form of the B-spline. Finally, these magenta points were fed to the informed autoencoder network in Fig. 2 to predict the open-close conformational transition of CRBN in full atomistic detail. The network predicted structures were then post-processed to relax some local geometric distortions (*22*) that are often found in ML-based structure predictions. A movie file and a Pymol session of the conformational transition are available in the Supplementary Materials.

**Figure 4:**
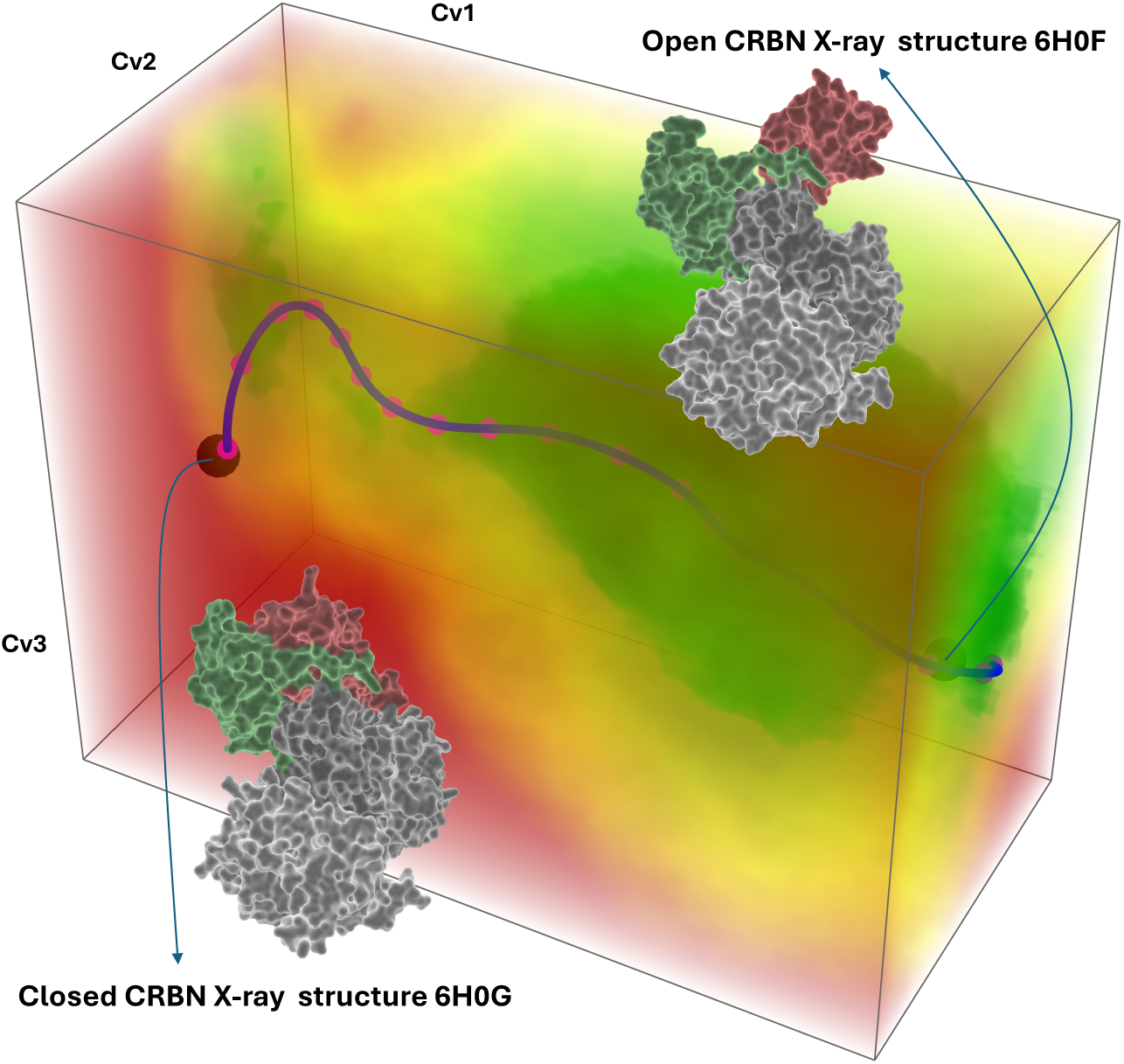
Putative low free-energy path of the CRBN open-close conformational transition. The free energy map shows that the volume of accessible closed conformations is very small compared to the very large number of feasible open conformations, and there is only a single narrow path connecting them. To establish a putative transition mechanism, we manually picked a few anchor points in the open, closed, and transition regions, respectively, then fitted a B-spline on them (blue curve), and computed the CV coordinates of the magenta points along the curve. These magenta points were then fed to the informed autoencoder network in Fig. 2 to predict the open-close conformational transition of CRBN in full atomistic detail. A movie file and a Pymol session are available in the Supplementary Materials. See main text and (22) for more details.

### Pros and cons of rMD

1. rMD is a simple and practical molecular simulation method that can add significant scientific value to an existing MD simulation with a trajectory and free energy map by quickly training an informed autoencoder network to generate new protein conformations, e.g. to model biologically relevant protein motions. Moreover, these new conformations can be used as seed structures for additional MD simulations to explore specific domains of the conformational space.
2. All trajectory frames from the MD simulation are superposed to a single frame, which is necessary to eliminate the global rotational and translational degrees of freedom before training the network (*22*). Since selection of the reference frame is arbitrary, there is straightforward data augmentation already built in in rMD. Multiple networks can be trained independently using different trajectory input data by simply superposing the structures on different randomly selected frames, respectively, and then using consensus to merge models generated from the same CV coordinates by different networks.
3. rMD can be easily used with the powerful concept of data reweighting in metadynamics simulations (*42–44*) where after computing a free energy map by biasing an MD simulation along one set of collective variables, one can reconstruct a different free energy map associated with another set of CVs, without performing additional simulation. rMD can then be used to quickly generate, e.g. alternative transition paths relevant to the new free energy map.
4. Conformational transition paths generated by rMD can be considered as high-quality initial guesses for more sophisticated path meta-eABF simulations to compute the free energy profile (potential of mean force) along the curvilinear transition path, including statistical error estimates (*45*).
5. Because rMD training requires the simultaneous optimization of two different loss functions, it is more difficult to lower the average reconstruction RMSD (Loss2) to close to the desired 1 Å. For example, in our network, the average Loss2 RMSD (including all heavy atoms) was ≈ 1.6 Å (*22*). We are currently working on alternative loss functions, for example, applying the Chamfer or Hausdorrf distances (*46*) for point clouds in Loss1 and including a local residue-based interatomic distance loss term (*7*) in Loss2.

## Acknowledgments

I. K. thanks Arben Kalziqi for help with the Wolfram language.

## Competing interests

There are no competing interests to declare.

## Data and materials availability

Wolfram code and simulation data will be available in the future.

## Supplementary materials

Materials and Methods

Figs. S1 to S2

References (*47–74*)

Movie S1

Data S1

## Materials and Methods

### Molecular dynamics simulations

We ran our molecular dynamics simulations using a combination of metadynamics and the extended Lagrangian adaptive biasing force (meta-eABF) method (*49, 50*) to bias the simulation along three collective variables (CV) and estimate the corresponding 3-dimensional free energy map. Meta-eABF simulations utilize an adaptive free-energy biasing force to enhance sampling along one or more CVs. Meta-eABF evokes the extended Lagrangian formalism of ABF whereby an auxiliary simulation is introduced with a small number of degrees of freedom equal to the number of CVs, and each real CV is associated with its so-called fictitious counterpart in the low-dimensional auxiliary simulation. The real CV is tied to its fictitious CV via a spring (typically with a large force constant), and the adaptive biasing force is equal to the running average of the negative of the spring force. The biasing force is only applied to the fictitious CV, which in turn “drags” the real simulation along the real CV via the spring by periodically injecting the instantaneous spring force back into the real simulation. Moreover, the main tenet of the meta-eABF method is that metadynamics is employed to enhance sampling of the fictitious CV itself. The combined approach has been shown to provide advantages over using metadynamics or eABF (*51*) alone.

We prepared two systems for simulation, one with the closed conformation (PDB ID 6H0G) and one with the open conformation (PDB ID 6H0F) of CRBN (*31, 32*). First, we ran two sets of exploratory simulations, one with the whole system and then another including only the DDB1-CRBN apo complex without Pomalidomide. Each set consisted of two 1 𝜇s long simulations starting from the CRBN-closed and CRBN-open conformations, respectively. The trajectories showed no discernible differences in terms of the open-closed CRBN conformational transition, and therefore, we chose the apo system for generating input data for the machine learning model. The missing residues were added with the Swiss-Model web server (*52*); followed by the preparation of the system with AmberTools (*53,54*) and the use of the force fields ff14sb (*55*), gaff (*56*), and the TIP3P water model (*57, 58*). We applied hydrogen mass repartitioning (HMR) (*59*), periodic boundary conditions with the PME (*60, 61*) long-range electrostatic model with a 9 Å real-space cutoff, in the NPT ensemble at 310 K and atmospheric pressure. We completed our simulations on a single Linux workstation equipped with two Nvidia RTX 4080 GPUs and 48 4-GHz AMD Ryzen CPU cores, running Ubuntu 22.04, and using the Openmm-Plumed software (*62–65*), versions 8.1.0 and 2.9.0, respectively. The primary simulation cell contained close to 210 000 atoms and the simulation wall-clock speed was 135 ns/day with a 4 fs time step facilitated by a rigid water model (*66*) and HMR with the mass of H atoms set at 3.024 amu (*59*).

### Collective variables

The meta-eABF bias was applied in a 3-dimensional collective variable (CV) space shown in Fig. S1 where we superposed the open and closed forms of the CRBN-DDB1 complex. The spheres represent the center of mass (COM) of the secondary structure regions of the domains CRBN-NTD (green), CRBN-CTD (red), CRBN-HBD (gray), and DDB1-BPC (teal), respectively. Since the open-close transition significantly displaces only the red COM, the CV space can be defined by the red sides CV1, CV2, and CV3 of the tetrahedron formed by the four COMs. Varying these three distances simultaneously is equivalent to the conical motion of the CRBN-CTD COM with respect to the base face of the tetrahedron. The conical motion of the red sphere inside the blue ellipsoid bubble explores a plethora of potential open-close transition mechanisms. The meta-eABF implementation in Plumed provides a (in our case 3-dimensional) grid of unbiased estimates of the free energy gradient using corrected Z-averaged constraints (CZAR) (*67*). We integrated the gradients with a Poisson integrator, which is part of the Colvars library (*68, 69*), to obtain the 3-dimensional free energy map.

### Informed autoencoder network

Figure S2 shows a detailed graph of our informed autoencoder network. We designed, trained and performed predictions with the network in a Wolfram/Mathematica 14.1 (*70*) notebook on the same desktop computer that we used for the MD simulations. The central piece of our network is the linked encoder-decoder chain with two different loss functions Loss1 and Loss2, respectively. On the top, the encoder is expanded to show all of its layers, and similarly, the decoder is detailed at the bottom. The basic design was inspired by (*9*) but we introduced an additional loss function to infuse physical meaning into the latent space. The encoder passes the flattened Cartesian coordinates of protein structures taken from trajectory frames of one or more MD simulations through a cascade of fully connected hidden layers with a gradually decreasing number of neurons. The final layer is called latent space (LS), where the original protein structures are compressed into a representation of only a few dimensions, in our case three dimensions. The decoder then takes the low-dimensional latent space coordinates as input and expands them back to the original Cartesian representation of protein conformations, using a similar set of now expanding hidden layers. The autoencoder design can be visualized as a butterfly with the thorax as the latent space. The autoencoder can be trained to optimize the reconstruction loss function—Loss2 in Fig. S2—such that the structural superposition between input and output is minimized, based on a set of training data.

The basic tenet of our autoencoder design is to introduce a new loss function–Loss1 in Fig. S2—that is applied to the latent space. In the original autoencoder, the latent space has no physical meaning, and although the LS coordinates can readily be used to generate new protein structures that are not encountered in the MD simulation(s), these structures are generic. Our informed autoencoder design empowers the network to perform targeted protein structure prediction by linking the latent space with the collective variable space. The input for the latent space loss function Loss1 are the LS coordinates and the target are the CV coordinates shown as the purple ”colVars” box. By simultaneously optimizing Loss1 and Loss2, e.g., using a weighted sum, the trained network will compress the original trajectory frames into a unique latent space where LS coordinates have an approximate one-to-one correspondence with the CV coordinates. This means that instead of using the otherwise meaningless LS coordinates for protein structure generation, we can use the 3-dimensional free-energy map directly since the map is defined over the CV space. This is how physics-based information is injected into the LS space.

The training data were the flattened Cartesian coordinates of all heavy atoms in the 10 000 trajectory frames from two independent meta-eABF simulations. Each simulation was 1 𝜇s long and frames were saved every 200 ps. In the simulations, the DDB1 protein did not show significant motion in addition to small thermal fluctuations, and therefore we only included CRBN in the training, resulting in length-9696 input vectors. It should be noted that, prior to training, all protein structures were superposed to the first frame, which means that using the mean absolute error for the Loss2 function was very close to the traditional root mean square distance (RMSD) widely used to measure structural superposition error. Each linear layer, except for the latent space layer in the encoder and the prediction layer in the decoder, was augmented with an element-wise activation layer using the ”Swish” form of the logistic sigmoid function 𝑥𝜎(𝑥). The training data were randomized and split into 8 000 structures for training and 2 000 structures for validation. We ran 10 000 training rounds with a batch size of 64 using the Adams optimizer. Training took 2 hours on the RTX 4080 GPU.

The final loss values of the trained network were Loss1 ≈ 1 Å and Loss2 ≈ 1.6 Å, which corresponds to ≈ 1.6 Å RMSD of all heavy atoms on average. It should be noted that although the network predicted generally high-quality structures, there were also numerous local distortions in the geometry, especially in flexible loops that had to be post-processed. This was of course expected since the network had no access to a dictionary of side-chain rotamers or other built-in knowledge base. We cleaned up the predicted structures in two steps. We used Rosetta Relax (*71–73*) on the Rosie server (*74*) followed by additional position-restrained minimization while keeping the C-alpha and C-beta atoms in place. The resulting final structures were virtually indistinguishable from those predicted by the network and all local distortions were fixed.

**Figure S1:**
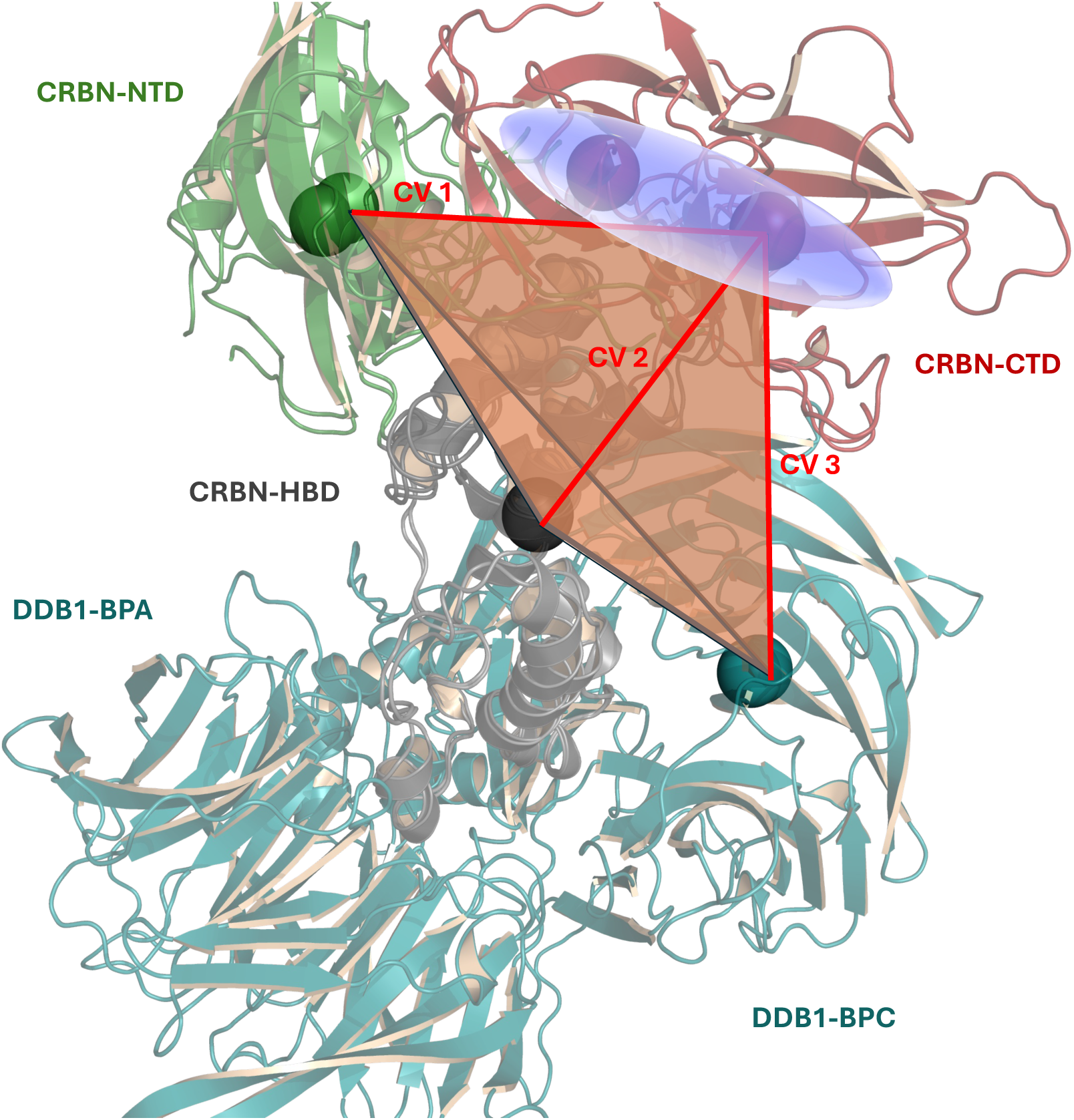
Definition of the collective variables. The spheres represent the center of mass (COM) of the secondary structure regions of the domains CRBN-NTD (green), CRBN-CTD (red), CRBN-HBD (gray), and DDB1-BPC (teal), respectively. Since the open-close transition significantly displaces only the red COM, the CV space can be defined by the red sides CV1, CV2, and CV3 of the tetrahedron. Varying these three distances simultaneously is equivalent to the conical motion of the red CRBN-CTD COM (inside the blue bubble) with respect to the base face of the tetrahedron. See (*22*) for more details.

**Figure S2:**
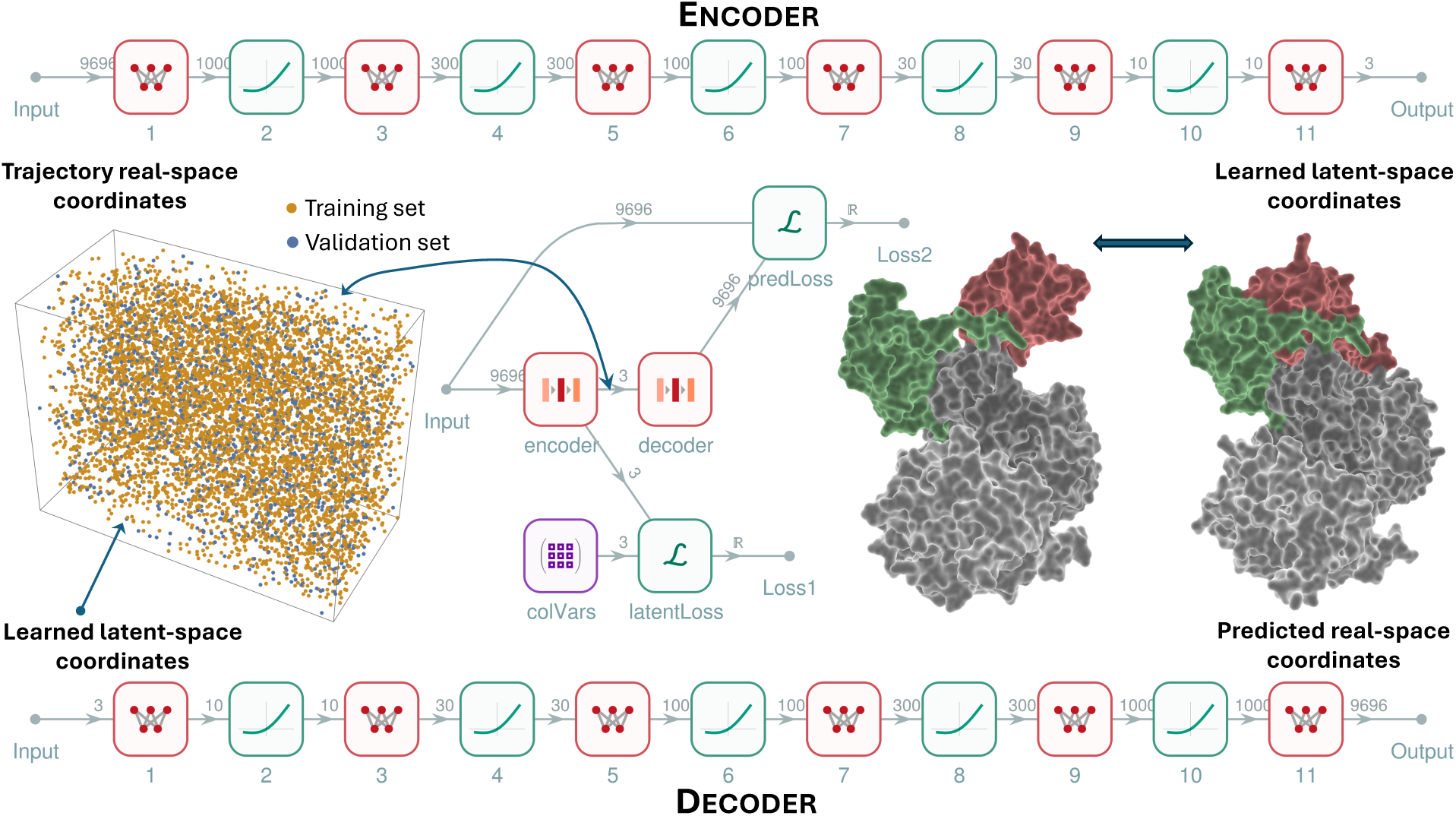
Net graph representation of the rMD informed autoencoder network. This is a more detailed version of Fig. 2 with the encoder and the decoder unfolded to show all the hidden layers and their dimensions. The encoder compresses the flattened Cartesian coordinates of protein structures from MD simulations into a low-dimensional latent space and the decoder reconstructs the original Cartesian representation from the LS coordinates. The rMD autoencoder is trained using two loss functions simultaneously. The ”predLoss” layer, minimizing the Loss2 function, which calculates the average RMSD between inputs and outputs, and the ”latentLoss” layer minimizing Loss1. The input for Loss1 are the LS coordinates and the target are the CV coordinates (purple ”colVars” box), and Loss1 computes the RMSD between the two point clouds. By simultaneously optimizing Loss1 and Loss2, the trained network will compress the original trajectory frames into a unique latent space where LS coordinates have an approximate one-to-one correspondence with the CV coordinates as shown in Fig. 3. Note that the optimized point distribution of the 10 000 input structures in LS space (training set orange, validation set blue) is the same as in Fig. 3.

**Caption for Movie S1. Animation of the open-close conformational transition of CRBN as predicted by the rMD autoencoder network.** This video shows the conformational transition along the blue B-spline curve in Fig. 4.

**Caption for Data S1. Pymol 3 session file used to create Movie S1.** The Pymol session contains two objects, each with 20 states that correspond to the network-predicted all-atom structures initiated from the magenta points in Fig. 4. The structures in the first object ”path” were post-processed as described in (*22*) and the structures in the other object ”path smooth” were further smoothed by Pymol for smooth animation.

## Notes

https://doi.org/10.5281/zenodo.14853955

https://doi.org/10.5281/zenodo.14853987

### Competing Interest Statement

The authors have declared no competing interest.

## References and Notes

1. J. Jumper, et al., Highly accurate protein structure prediction with AlphaFold. Nature 596 (7873), 583–589 (2021).

2. J. Abramson, et al., Accurate structure prediction of biomolecular interactions with AlphaFold Nature pp. 1–3 (2024).

3. M. Baek, et al., Accurate prediction of protein structures and interactions using a three-track neural network. Science 373 (6557), 871–876 (2021).

4. G. Ahdritz, et al., OpenFold: Retraining AlphaFold2 yields new insights into its learning mechanisms and capacity for generalization. Nature Methods pp. 1–11 (2024).

5. Z. Lin, et al., Evolutionary-scale prediction of atomic-level protein structure with a language model. Science 379 (6637), 1123–1130 (2023).

6. K. E. Wu, et al., Protein structure generation via folding diffusion. Nature communications 15 (1), 1059 (2024).

7. A. Liu, A. Elaldi, N. Russell, O. Viessmann, Bio2Token: All-atom tokenization of any biomolecular structure with Mamba. arXiv (2024), arXiv:2410.19110.

8. S. Lewis, et al., Scalable emulation of protein equilibrium ensembles with generative deep learning. bioRxiv pp. 2024–12 (2024), bioRxiv:2024.12.05.626885.

9. M. T. Degiacomi, Coupling molecular dynamics and deep learning to mine protein conformational space. Structure 27 (6), 1034–1040 (2019).

10. Y. Jin, L. O. Johannissen, S. Hay, Predicting new protein conformations from molecular dynamics simulation conformational landscapes and machine learning. Proteins: structure, function, and bioinformatics 89 (8), 915–921 (2021).

11. M. Ghorbani, S. Prasad, J. B. Klauda, B. R. Brooks, Variational embedding of protein folding simulations using Gaussian mixture variational autoencoders. The Journal of Chemical Physics 155 (19) (2021).

12. H. Tian, et al., Explore protein conformational space with variational autoencoder. Frontiers in molecular biosciences 8, 781635 (2021).

13. S. Xiao, Z. Song, H. Tian, P. Tao, Assessments of Variational Autoencoder in Protein Conformation Exploration. Journal of computational biophysics and chemistry 22 (04), 489–501 (2023).

14. S. Mansoor, M. Baek, H. Park, G. R. Lee, D. Baker, Protein Ensemble Generation through Variational Autoencoder Latent Space Sampling. Journal of Chemical Theory and Computation 20 (7), 2689–2695 (2024).

15. B. P. Vani, A. Aranganathan, D. Wang, P. Tiwary, Alphafold2-rave: From sequence to boltzmann ranking. Journal of chemical theory and computation 19 (14), 4351–4354 (2023).

16. D. Teng, V. J. Meraz, A. Aranganathan, X. Gu, P. Tiwary, AlphaFold2-RAVE: Protein Ensemble Generation with Physics-Based Sampling. chemRxiv (2025), chemrxiv:2025-q3mwr.

17. V. K. Ramaswamy, S. C. Musson, C. G. Willcocks, M. T. Degiacomi, Deep learning protein conformational space with convolutions and latent interpolations. Physical Review X 11 (1), 011052 (2021).

18. M. D. Ward, et al., Deep learning the structural determinants of protein biochemical properties by comparing structural ensembles with DiffNets. Nature communications 12 (1), 3023 (2021).

19. S. Kaptan, I. Vattulainen, Machine learning in the analysis of biomolecular simulations. Advances in physics: X 7 (1), 2006080 (2022).

20. L.-E. Zheng, S. Barethiya, E. Nordquist, J. Chen, Machine learning generation of dynamic protein conformational ensembles. Molecules 28 (10), 4047 (2023).

21. M. H. Murtada, Z. F. Brotzakis, M. Vendruscolo, Language Models for Molecular Dynamics. bioRxiv pp. 2024–11 (2024), bioRxiv:2024.11.25.625337.

22. Materials and methods are available as supplementary material.

23. J. B. Bartlett, K. Dredge, A. G. Dalgleish, The evolution of thalidomide and its IMiD derivatives as anticancer agents. Nature Reviews Cancer 4 (4), 314–322 (2004), doi:10.1038/nrc1323.

24. T. Ito, et al., Identification of a Primary Target of Thalidomide Teratogenicity. Science 327 (5971), 1345–1350 (2010), doi:10.1126/science.1177319.

25. J. Krönke, et al., Lenalidomide Causes Selective Degradation of IKZF1 and IKZF3 in Multiple Myeloma Cells. Science 343 (6168), 301–305 (2014), doi:10.1126/science.1244851.

26. J. M. Sasso, et al., Molecular Glues: The Adhesive Connecting Targeted Protein Degradation to the Clinic. Biochemistry 62 (3), 601–623 (2022), doi:10.1021/acs.biochem.2c00245.

27. M. F. Neurath, L. J. Berg, VAV1 as a putative therapeutic target in autoimmune and chronic inflammatory diseases. Trends in Immunology 45 (8), 580–596 (2024), doi:10.1016/j.it.2024.06.004.

28. A. Sylvain, et al., A cereblon (CRBN) molecular glue degrader of NIMA-related kinase 7 (NEK7) reveals a context-dependent role in NLRP3 inflammasome activation. bioRxiv (2024), bioRxiv:2024.11.06.622079v1, doi:10.1101/2024.11.06.622079.

29. G. Nishiguchi, et al., Selective CK1𝛼 degraders exert antiproliferative activity against a broad range of human cancer cell lines. Nature Communications 15 (1) (2024), doi: 10.1038/s41467-024-44698-1.

30. G. Petzold, E. S. Fischer, N. H. Thomä, Structural basis of lenalidomide-induced CK1𝛼 degradation by the CRL4CRBN ubiquitin ligase. Nature 532 (7597), 127–130 (2016), doi: 10.1038/nature16979.

31. Q. L. Sievers, et al., Defining the human C2H2 zinc finger degrome targeted by thalidomide analogs through CRBN. Science 362 (6414) (2018), doi:10.1126/science.aat0572.

32. E. R. Watson, et al., Molecular glue CELMoD compounds are regulators of cereblon conformation. Science 378 (6619), 549–553 (2022), doi:10.1126/science.add7574.

33. A. Laio, M. Parrinello, Escaping free-energy minima. Proceedings of the national academy of sciences 99 (20), 12562–12566 (2002).

34. G. Bussi, F. L. Gervasio, A. Laio, M. Parrinello, Free-energy landscape for 𝛽 hairpin folding from combined parallel tempering and metadynamics. Journal of the American Chemical Society 128 (41), 13435–13441 (2006).

35. S. Piana, A. Laio, A bias-exchange approach to protein folding. The journal of physical chemistry B 111 (17), 4553–4559 (2007).

36. A. Barducci, G. Bussi, M. Parrinello, Well-tempered metadynamics: a smoothly converging and tunable free-energy method. Physical review letters 100 (2), 020603 (2008).

37. P. Tiwary, M. Parrinello, From metadynamics to dynamics. Physical review letters 111 (23), 230602 (2013).

38. M. Invernizzi, M. Parrinello, Rethinking metadynamics: from bias potentials to probability distributions. The journal of physical chemistry letters 11 (7), 2731–2736 (2020).

39. V. Rizzi, S. Aureli, N. Ansari, F. L. Gervasio, OneOPES, a combined enhanced sampling method to rule them all. Journal of Chemical Theory and Computation 19 (17), 5731–5742 (2023).

40. P. Izmailov, D. Podoprikhin, T. Garipov, D. Vetrov, A. G. Wilson, Averaging weights leads to wider optima and better generalization. arXiv (2018), arXiv:1803.05407.

41. Ideally, we would plot the locations of the apo open and closed CRBN X-ray structures, but in lieu of that PDB IDs 6H0G and 6H0F are the closest match.

42. D. Branduardi, G. Bussi, M. Parrinello, Metadynamics with adaptive Gaussians. Journal of chemical theory and computation 8 (7), 2247–2254 (2012).

43. P. Tiwary, M. Parrinello, A time-independent free energy estimator for metadynamics. The Journal of Physical Chemistry B 119 (3), 736–742 (2015).

44. J. Rydzewski, M. Chen, T. K. Ghosh, O. Valsson, Reweighted manifold learning of collective variables from enhanced sampling simulations. Journal of Chemical Theory and Computation 18 (12), 7179–7192 (2022).

45. I. Kolossváry, W. Sherman, Comprehensive Approach to Simulating Large Scale Conformational Changes in Biological Systems Utilizing a Path Collective Variable and New Barrier Restraint. The Journal of Physical Chemistry B 127 (23), 5214–5229 (2023).

46. T. Nguyen, et al., Point-set distances for learning representations of 3d point clouds, in *Proceedings of the IEEE/CVF international conference on computer vision* (2021), pp. 10478–10487.

47. I. Kolossváry, ML-generated CRBN open-closed transition path, Zenodo (2025), doi:10.5281/ zenodo.14853955, 10.5281/zenodo.14853955.

48. I. Kolossváry, Pymol session of the ML simulated CRBN open-closed conformational transition, Zenodo (2025), doi:10.5281/zenodo.14853987, 10.5281/zenodo.14853987.

49. J. Comer, et al., The adaptive biasing force method: Everything you always wanted to know but were afraid to ask. J. Phys. Chem. B 119 (3), 1129–1151 (2015).

50. H. Fu, et al., Zooming across the free-energy landscape: shaving barriers, and flooding valleys. J. Phys. Chem. Lett. 9 (16), 4738–4745 (2018).

51. H. Fu, X. Shao, W. Cai, C. Chipot, Taming rugged free energy landscapes using an average force. Acc. Chem. Res. 52 (11), 3254–3264 (2019).

52. T. Schwede, SWISS-MODEL, SWISS-MODEL, https://swissmodel.expasy.org/.

53. D. A. Case, et al., The Amber biomolecular simulation programs. Journal of computational chemistry 26 (16), 1668–1688 (2005).

54. R. Salomon-Ferrer, D. A. Case, R. C. Walker, An overview of the Amber biomolecular simulation package. Wiley Interdisciplinary Reviews: Computational Molecular Science 3 (2), 198–210 (2013).

55. J. A. Maier, et al., ff14SB: improving the accuracy of protein side chain and backbone parameters from ff99SB. Journal of chemical theory and computation 11 (8), 3696–3713 (2015).

56. J. Wang, R. M. Wolf, J. W. Caldwell, P. A. Kollman, D. A. Case, Development and testing of a general amber force field. Journal of computational chemistry 25 (9), 1157–1174 (2004).

57. W. L. Jorgensen, J. Chandrasekhar, J. D. Madura, R. W. Impey, M. L. Klein, Comparison of simple potential functions for simulating liquid water. The Journal of chemical physics 79 (2), 926–935 (1983).

58. D. J. Price, C. L. Brooks III, A modified TIP3P water potential for simulation with Ewald summation. The Journal of chemical physics 121 (20), 10096–10103 (2004).

59. C. W. Hopkins, S. Le Grand, R. C. Walker, A. E. Roitberg, Long-time-step molecular dynamics through hydrogen mass repartitioning. Journal of chemical theory and computation 11 (4), 1864–1874 (2015).

60. T. Darden, D. York, L. Pedersen, Particle mesh Ewald: An N log (N) method for Ewald sums in large systems. The Journal of chemical physics 98 (12), 10089–10092 (1993).

61. U. Essmann, et al., A smooth particle mesh Ewald method. The Journal of chemical physics 103 (19), 8577–8593 (1995).

62. P. Eastman, et al., OpenMM 7: Rapid development of high performance algorithms for molecular dynamics. PLoS Comput. Biol. 13 (7), e1005659 (2017).

63. M. Bonomi, Promoting transparency and reproducibility in enhanced molecular simulations. Nat. Methods 16 (8), 670–673 (2019).

64. M. Bonomi, et al., PLUMED: A portable plugin for free-energy calculations with molecular dynamics. Comput. Phys. Commun. 180 (10), 1961–1972 (2009), 10.1016/j.cpc.2009.05.011, https://www.sciencedirect.com/science/article/pii/S001046550900157X.

65. G. A. Tribello, M. Bonomi, D. Branduardi, C. Camilloni, G. Bussi, PLUMED 2: New feathers for an old bird. Comput. Phys. Commun. 185 (2), 604–613 (2014), 10.1016/j.cpc.2013.09.018, https://www.sciencedirect.com/science/article/pii/S0010465513003196.

66. S. Miyamoto, P. A. Kollman, Settle: An analytical version of the SHAKE and RATTLE algorithm for rigid water models. Journal of computational chemistry 13 (8), 952–962 (1992).

67. A. Lesage, T. Lelievre, G. Stoltz, J. Hénin, Smoothed biasing forces yield unbiased free energies with the extended-system adaptive biasing force method. J. Phys. Chem. B 121 (15), 3676–3685 (2017).

68. G. Fiorin, M. L. Klein, J. Hénin, Using collective variables to drive molecular dynamics simulations. Mol. Phys. 111 (22-23), 3345–3362 (2013).

69. G. Fiorin, et al., Expanded functionality and portability for the Colvars library. The Journal of Physical Chemistry B 128 (45), 11108–11123 (2024).

70. Wolfram Research, Inc., Mathematica (2024), https://www.wolfram.com/mathematica.

71. M. D. Tyka, et al., Alternate states of proteins revealed by detailed energy landscape mapping. Journal of molecular biology 405 (2), 607–618 (2011).

72. F. Khatib, et al., Algorithm discovery by protein folding game players. Proceedings of the National Academy of Sciences 108 (47), 18949–18953 (2011).

73. J. B. Maguire, et al., Perturbing the energy landscape for improved packing during computational protein design. Proteins: Structure, Function, and Bioinformatics 89 (4), 436–449 (2021).

74. S. Lyskov, et al., Serverification of molecular modeling applications: the Rosetta Online Server that Includes Everyone (ROSIE). PloS one 8 (5), e63906 (2013).

